# TransMEP: Transfer learning on large protein language models to predict mutation effects of proteins from a small known dataset

**DOI:** 10.1101/2024.01.12.575432

**Authors:** Tilman Hoffbauer, Birgit Strodel

**Affiliations:** Institute of Biological Information Processing: Structural Biochemistry (IBI-7), Forschungszentrum Jülich, 52428 Jülich, Germany; RWTH Aachen University, 52062 Aachen, Germany; Institute of Theoretical and Computational Chemistry, Heinrich Heine University Düsseldorf, 40225 Düsseldorf, Germany

**Keywords:** machine learning, transfer learning, protein language model, Gaussian process, upper confidence bound optimization, protein engineering, mutation effect prediction

## Abstract

Machine learning-guided optimization has become a driving force for recent improvements in protein engineering. In addition, new protein language models are learning the grammar of evolutionarily occurring sequences at large scales. This work combines both approaches to make predictions about mutational effects that support protein engineering. To this end, an easy-to-use software tool called TransMEP is developed using transfer learning by feature extraction with Gaussian process regression. A large collection of datasets is used to evaluate its quality, which scales with the size of the training set, and to show its improvements over previous fine-tuning approaches. Wet-lab studies are simulated to evaluate the use of mutation effect prediction models for protein engineering. This showed that TransMEP finds the best performing mutants with a limited study budget by considering the trade-off between exploration and exploitation.

**Graphical TOC Entry:** 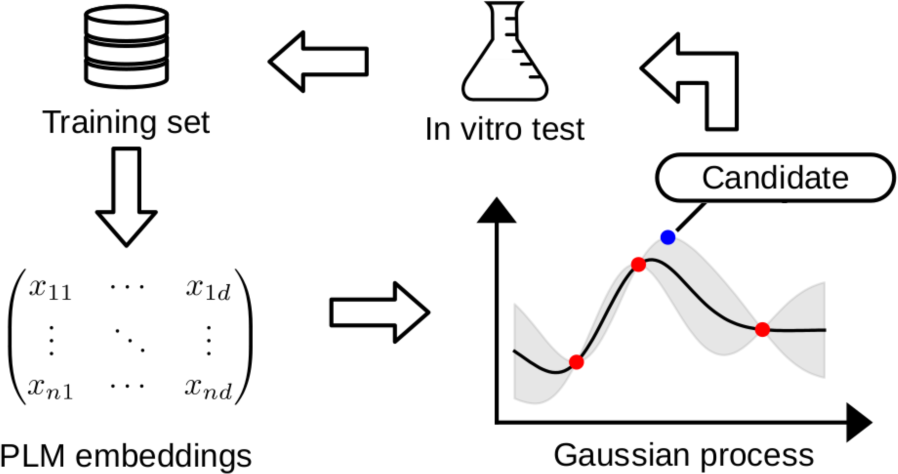

## 1 Introduction

Protein engineering involves optimizing a given amino acid sequence for a user-specified target by introducing mutations into the sequence that alter the protein’s properties. ^1^ It is particularly applied to enzymes to increase their stability, activity, or enantioselectivity so that they can be used in industrial processes.^2,3^ However, selecting promising mutations to tune the property of interest based on current knowledge, so-called rational design, is often difficult due to unpredictable epistatic effects.^4^ An important approach in this context is the so-called directed evolution.^5^ The inherent bottleneck of this approach is the cost associated with the production and evaluation of the of the huge mutant library.^6^ Therefore, it is advantageous to carefully select the mutants to be tested. Machine learning has recently gained traction as an in silico tool to predict the expected effect of mutations based on known in vitro assessed mutants.^7–12^

A machine learning approach to predicting mutational effects usually consists of two main components: First, features, i.e., a set of numerical values, are calculated for each mutant; second, a machine learning model is fitted to these features to predict the desired target value, such as the stability or activity of an enzyme. Several commonly used transformations are available for the features. As a naive approach, one can encode the amino acid sequence of a protein by describing each residue with a one-hot vector corresponding to the amino acid at that position. Alternatively, each amino acid can be represented by a set of chemical descriptors, such as the polarity of an amino acid. This can be extended by contextual embeddings, where the features of a residue depend not only on the amino acid but also on neighboring residues. Many of these descriptors have been compared by Xu et al. ^8^ Protein language models (PLMs) ^9,13,14^ take this idea even further. In this approach, a large neural network is pre-trained on millions of sequences of proteins with unknown functions and structures to predict a masked amino acid from its context. This forces the model to learn statistical relationships between amino acids in naturally occurring sequences. The derived model can then be used as a foundation model for generating contextual embeddings used as input for downstream models, which is known as transfer learning via feature extraction.^15^ It was shown that these descriptors improve predictions over existing machine learning approaches on a wide variety of proteins and problems. ^9,16^ The current state-of-theart of these PLMs was published by Lin et al. from Meta AI Research (former Facebook AI Research) as ESM-2 (Evolutionary Scale Modeling 2). ^14^

For model selection, the designer is confronted with many different options. Many of these have been compared by Xu et al. and Yang et al. who proposed a model selection flow chart.^7,8^ To select the best model automatically, Wittmann et al. created a tool that fits multiple models simultaneously and then compares the cross-validation error to build an ensemble of the top-performing models.^10^ Rives et al. used a different approach.^9^ They replaced the final layer of ESM with a new classification head and fine-tuned the whole model on various datasets with gradient descent. To apply a mutation effect prediction model to a protein engineering problem, one must solve the additional problem of selecting candidates to be tested next in the wet lab. Wittmann et al. proposed a method that always samples the next candidates with the best target value as predicted by the current model.^10^ Importantly, they have shown that careful selection of the variants to be sampled is critical to the accuracy of the method. In the approach of Greenhalgh et al., a Gaussian process model was used that simultaneously predicts the mean and standard deviation of the target value.^11^ This allows the application of a well-known optimization scheme, the so-called upper confidence bound optimization, an optimization method with sound theoretical support.^17^ In upper confidence optimization applied to protein engineering, the user can control the trade-off between the exploration of the mutant space and the exploitation of interesting candidates.

Here we present our machine learning approach for protein engineering called TransMEP (Transfer learning for Mutation Effect Prediction), an implementation of the transfer learning approach that is easy to use, fast and simple to apply. Unlike other work, TransMEP combines the use of a large transfer learning model and the simulation of mutagenesis studies, taking into account the trade-off between exploration and exploitation. This is enabled by the high performance of the model, which is achieved through the efficient use of a GPU accelerator and other optimizations. In addition, we provide a thorough comparison between fine-tuning and using fixed embeddings. We demonstrate the good predictive quality of TransMEP for various datasets and simulate wet-lab protein engineering studies with it.

## 2 Methodology

### 2.1 Protein language models

In this work, the 150 million parameters variant of ESM-2 (esm2 t30 150M UR50D)^14^ is used, although TransMEP can be configured to use other variants of ESM. This model is a variant of the BERT (Bidirectional Encoder Representations from Transformers)^18^ language model trained on raw amino acid sequences of naturally occurring proteins from the UniRef database.^19^ During training, 15% of the residues are masked and the model is tasked with reproducing the original amino acids from the remaining sequence and the knowledge acquired during training. To do this, it learns an internal representation for each residue, called embedding, that depends on the other residues. In fact, an embedding is most influenced by the structurally nearby residues, i.e., the PLM learns a partial solution to the folding problem. As such, the embeddings of PLMs capture interesting properties useful for downstream tasks, as, for instance, demonstrated by Rives et al. when fine-tuning the first version of ESM to predict mutation effects.^9^ The embeddings can be interpreted as a representation of the proteins in a high-dimensional space with a euclidean metric, as experiments by Elnaggar et al. showed.^13^ For further explanations on language models, the interested reader is referred to Chapters 11 and 15 of the book by Zhang et al. ^20^ which provides a tutorial on these topics. In TransMEP, the concatenation of all residue embeddings is used for the embedding space *X* = R*^d^* where *d ∈* N is the dimensionality of these embeddings, which is influenced by both the design of the PLM and the sequence length. Although this slightly abuses notation, we define set *X ^n^* = R*^n×d^*, where the *n* samples are arranged along the rows and not along the columns of the matrix.

### 2.2 Gaussian process regression

The introduction to Gaussian processes presented here is based on Chapter 2 of the book by Rasmussen and Williams,^21^ to which the interested reader is referred for more information than is presented below. Gaussian proess regression models the target value function *f* : *X →* R as being observed from a Gaussian process, i.e., for all finite sets of query points **x** *∈ X ^k^, k ∈* N, we assume

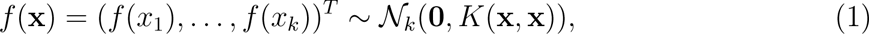

where *N_k_* is the *k*-variate normal distribution and *K*(*·, ·*) is the covariance matrix. In this work, the covariance matrix is expressed by a normalized radial basis function (RBF) kernel. The kernel function defines the covariance between two input sets **x** *∈ X ^k^* and **x***^i^ ∈ X ^l^* with lengths *k, l ∈* N as

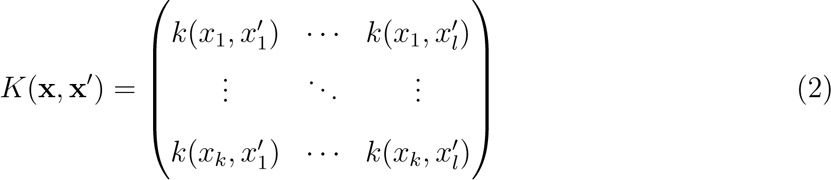

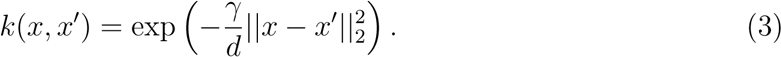

Note that *K*(**x***^i^,* **x**) = *K*(**x**, **x***^i^*)*^T^* . This kernel enforces the assumption that the target values of nearby embeddings *x, x^i^ ∈ X* are highly correlated, i.e. that the target value functions *f* (*x*) are smooth on a length scale *γ ∈* R. The length scale is normalized by *d* to reduce the changes in *γ* induced by varying embedding dimensionalities.

In the regression context, one is provided with a set of training data points **x** *∈ X ^n^* and query data points **x***^∗^ ∈ X ^m^* where *n, m ∈* N. Noisy observations are available for the training data points **y** = *f* (**x**) + ***E*** *∈* R*^n^* with ***E*** *∼ N_n_*(**0***, αI_n_*)*, α >* 0 being independent and identically distributed Gaussian noise. Following a Bayesian setting, this allows for modeling the joint distribution of noisy observations **y** and the target values of the query data points *f* (**x***^∗^*) as

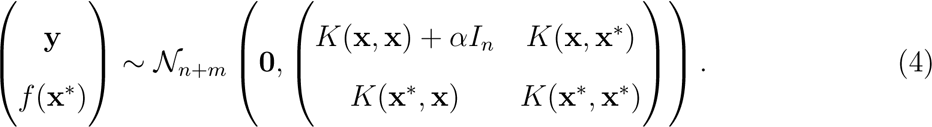

Here, *I_n_* denotes the *n*-dimensional identity matrix. As the conditional of a multivariate normal distribution is also multivariate normal distributed, and **y** is observed, this leads to

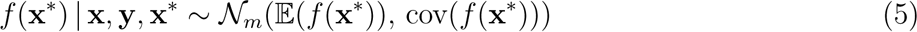

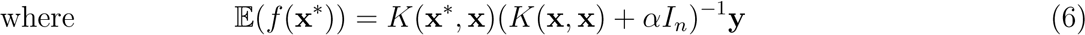

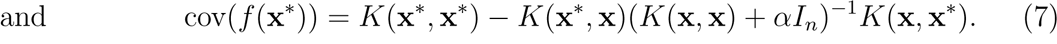

The choice of Gaussian process regression has several advantages over other machine learning algorithms. First, we not only get an estimate of the target value but a distribution of it, which allows for upper confidence bound optimization. Second, the use of a kernel function overcomes the problem of the high dimensionality of the embedding space *X* as the dimensions are coerced into one value by the euclidean norm. Third, it only has two hyper-parameters (*α, γ*) that control the noise and smoothness of the target values. Fourth, our previous work has shown that kernel ridge regression with an RBF kernel, which is a special case of Gaussian process regression with the same kernel,^21^ is well-suited for the application on PLM embeddings.^22^

### 2.3 Hyperparameter optimization

TransMEP uses grid search for automatic hyperparameter optimization (HPO) during each fit to a new dataset. For this, all combinations of *α* and *γ* on a 50 *×* 50 grid with *α* ranging from 10*^−^*^4^ to 10^5^ and *γ* ranging from 10*^−^*^3^ to 10^6^, both on a logarithmic scale, are evaluated. Each combination is tested on a randomly sampled set of 1,000 splits of the dataset into a training and a validation dataset to select the most promising hyperparameter values by the lowest mean squared error. This leads to a total number of 2,500,000 model fits, which is only possible due to the high parallelization achievable on GPUs and the small dataset sizes (at most 5,000 samples) one finds in typical protein engineering studies.

The HPO process benefits from the following important optimizations: First, the euclidean distances between all embeddings can be pre-computed, effectively eliminating the dependency on the high dimensionality of the embeddings and providing an important speedup. Second, the same splits are used for all hyperparameter values, which enables the alignment of these splits into memory-contiguous arrays a priori, allowing for batch processing of multiple splits simultaneously. Third, the Cholesky decomposition used for implicitly calculating the kernel matrix inverse (see Supporting Information for details) can be efficiently parallelized by cuSOLVER on NVIDIA GPUs.^23^ Fourth, the use of PyTorch^24^ allows for a simple-to-read, but very fast implementation of this process that can fall back to CPUs if no GPUs are available. TransMEP uses PyTorch and cuSOLVER for GPU acceleration of inference on Gaussian processes, too.

### 2.4 Upper confidence bound optimization

To select a good sample from the current model to test next, one must find a compromise between two opposing goals. On the one hand, the exploration of the mutation space is important to find good samples. On the other hand, the exploitation of predicted promising candidates is key to optimizing the target value. As such, neither pure exploration, which would be equivalent to random search, nor pure exploitation, which would be equivalent to just using the best prediction, is optimal. This is a well-known issue in optimization problems.^17^

Similar to the work by Greenhalgh et al.,^11^ we use the upper confidence bound (UCB) criterion to achieve this trade-off. The UCB criterion is defined as

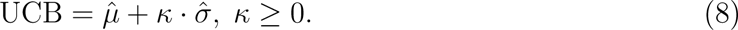

for a maximization problem, where *µ*^ is the predicted mean, *σ*^ the predicted standard deviation, and *κ* a hyperparameter. As such, it can only be applied to models that predict both a mean and a standard deviation, like Gaussian process regression. In each step, the next sample is chosen such that it maximizes this criterion and the model is updated with the new training data. Here, lower *κ* values correspond to more exploitation, while higher values of *κ* correspond to more exploration.^17^

### 2.5 Sampling candidates with high UCB values

As the UCB criterion is dependent on the prediction and not on the sequence, but in vitro testing requires the latter, a scheme to find sequences with high UCB values is required. In a naive approach, one might sample random mutants, pass them through the model and compute their UCB value. After several iterations, one would select the best mutant found. This brute-force approach suffers from the large search space, as the number of possible mutants grows exponentially with the number of allowed mutations. To improve over this naive approach, a genetic algorithm based on DEAP (Distributed Evolutionary Algorithms in Python)^25^ with an elitist principle as in NSGA-II^26^ is employed in TransMEP. Genetic algorithms are inspired by the evolutionary process and iteratively optimize a population of *n* individuals by crossover, mutation and selection. In this approach, mutants *x* are represented by a tuple of *k* mutations *x_i_* = (*p, a*)*, i ∈ {*1*, . . . , k}*, where *p* is the position and *a* the amino acid. All other residues are assumed to be identical to the wild-type protein. This ensures that each mutant may have at most *k* mutations at once, allowing the user to easily specify the maximum number of mutations to be considered. During optimization, the following scheme is iterated until convergence or a specified number of maximum iterations *m* is reached:

1. **Tournament selection:** Randomly sample two mutants and select the one with a higher UCB value *n* times, creating an offspring population of *n* individuals.
2. **Crossover:** Group the offspring population into pairs of two mutants *x, x^i^* and randomly exchange mutations *x_i_, x^i^* with probability *p_c_* for each index *i ∈ {*1*, . . . , k}*. Skip an index if this would lead to a mutant with two mutations for the same position.
3. **Mutation:** Randomly replace *x_i_* by a random mutation with probability *p_m_* for each mutant *x* in the offspring and index *i ∈ {*1*, . . . , k}*. Again, skip an index if this would lead to an invalid mutant.
4. **Elitist selection:** Select the *n* mutants with highest UCB value from both the previous population and the offspring.

Here, the population size *n*, crossover probability *p_c_*, mutation probability *p_m_*, and maximum number of iterations are hyperparameters. As the elitist selection principle creates populations where some mutants are repeated over multiple iterations, all computed UCB values are cached during optimization. This approach can trivially be extended to a criterion other than UCB.

### 2.6 Application of TransMEP

TransMEP is available at https://github.com/strodel-group/TransMEP where also the installation and usage of the software is described. For applying TransMEP to a given protein, one must be able to measure the desired fitness value (e.g., fluorescence or enantioselectivity) of about 100 mutants in total in the wet lab. To start the study, TransMEP should be provided with an initial set of approximately 10 known mutant/fitness pairs. Based on these, the user can employ TransMEP to generate new recommendations for protein mutations that should be tested next in vitro. This includes calculating the embeddings, fitting the model, and optimizing the UCB criterion, which usually takes about an hour. In the wet lab, the target values for the suggested mutants is determined, which is then fed back to TransMEP to generate the next mutant suggestions. This cycle is repeated until a sufficiently high target value is achieved, which should be the case after the evaluation of *∼*100 mutants. Further details on the usage of TransMEP can be found in the Supporting Information. In particular, two procedures are explained there that enable the interpretability of the models generated by TransMEP. One of them allows to determine the importance of the different training sets for mutation prediction, while another procedure facilitates the determination of the effect of a single mutation on the protein variant with multiple mutations.

## 3 Results

To verify the methods introduced above, we performed three computational experiments. First, the prediction accuracy of TransMEP is compared to the alternative approach of fine-tuning a PLM. Second, wet-lab studies are simulated to assess if our approach is able to find good mutants with a limited amount of mutant evaluations. Third, the sampling process is compared to random search and the mutants from a given, large dataset to show its effectiveness in finding mutants with a high UCB value.

### 3.1 Comparison to fine-tuning

#### Setup

TransMEP uses a fixed PLM, while other approaches often use fine-tuning^9^ as discussed in the introduction. We therefore compare these two methods in terms of prediction quality. To this end, we fine-tuned ESM-2^14^ on each dataset split by replacing the final amino acid prediction layer with a mean aggregation followed by dropout^28^ (*p* = 0.6) and a linear layer. The model was fine-tuned using the Adam optimizer^29^ with a learning rate of 10*^−^*^5^ on 90% of the training data, while the prediction quality on the remaining 10% of the data was monitored. As soon as this quantity did not improve for 50 epochs, the training was stopped and the model was reset to the best known previous state. To improve the initialization of the new linear layer, it was pre-trained for 10 epochs separately before unfreezing the full ESM-2 model. As fine-tuning is significantly slower than TransMEP on small datasets, the available computational budget limited the HPO to some hand-tuning steps which reduces its quality.

#### Data

Riesselman et al. ^27^ assembled a large collection of various protein engineering datasets. The proteins considered differ in function, organism from which they originate, size, and structure. For this computational experiment, each dataset was split into multiple training and test sets with training set sizes ranging from 50 to 5, 000, limited by the actual dataset size. For each size, five different splits were performed to increase measurement precision.

#### Results

In general, prediction quality improves with training set size (Figure 1). On average, fine-tuning performs significantly worse than TransMEP, while also having more variance. The performance gap often reduces with increasing training set size, and sometimes finetuning takes the lead. However, there are also some datasets where fine-tuning does not improve over the considered training set sizes.

**Figure 1:**
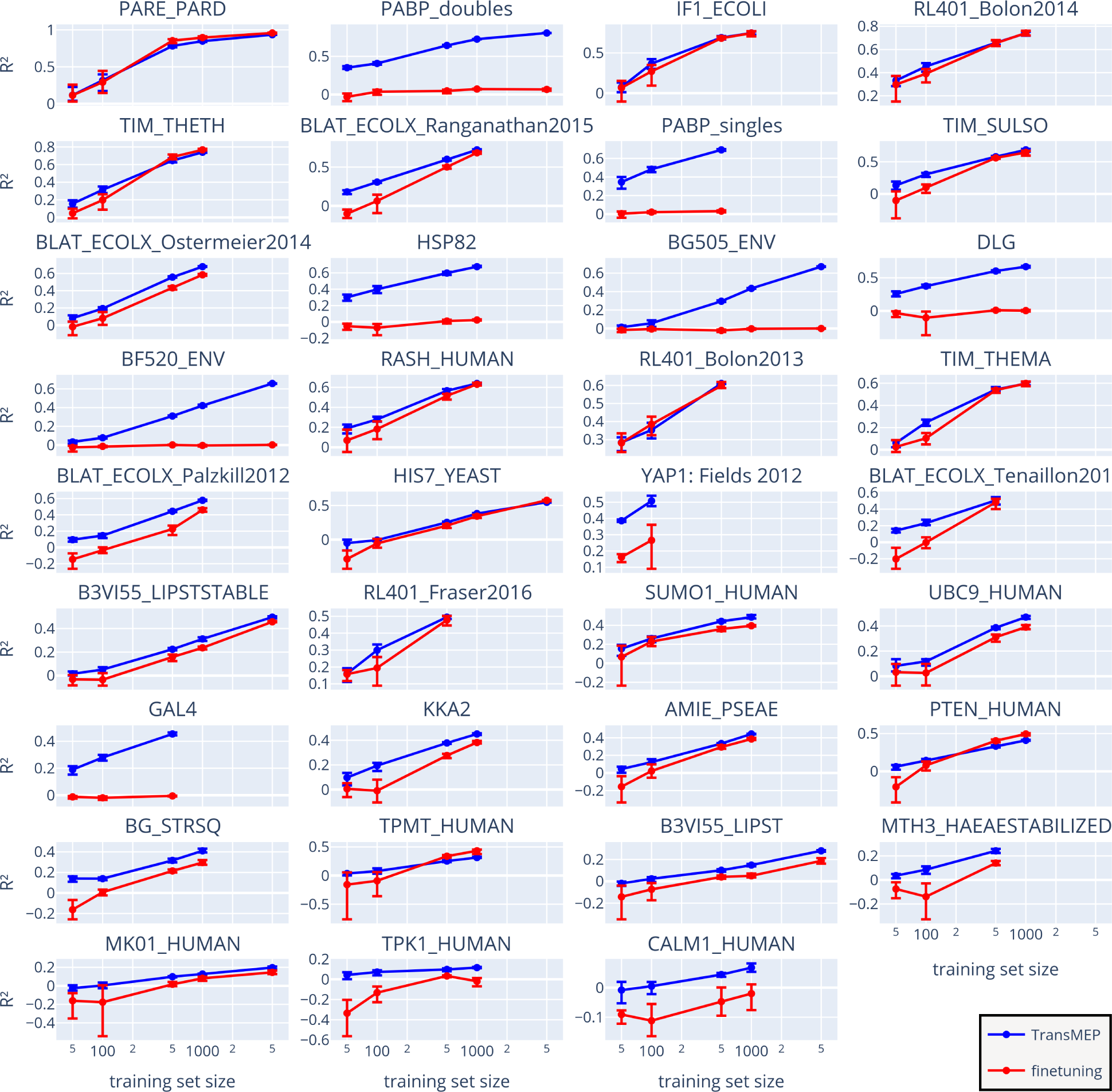
Performance of TransMEP (blue) and fine-tuning (red) measured as the fraction of explained variance (R^2^) for various datasets from Riesselman et al. ^27^ and different training set sizes (displayed on a logarithmic scale). Each measurement is shown as the mean of five different dataset splits and with the corresponding 90% bootstrap confidence interval. Note the different *y*-axis scales for the different data sets, which were chosen based on the minimum and maximum R^2^ values.

#### Discussion

The overall prediction quality is often not satisfying for either model, especially not for small training set sizes. The main drawback of this comparison is the lack of a thorough HPO for each fine-tuning run, similar to the procedure used by TransMEP. This can be seen as a major advantage of the latter approach: Since the PLM remains static, the embeddings and pairwise distances can be precomputed, and no gradient of the PLM is required. This saves computing time that can instead be used for a thorough HPO and additional studies thereafter. With TransMEP, there are only few runs that resulted in a negative coefficient of determination, i.e., a prediction worse than the mean of the test set, while fine-tuning more often fell below this baseline. These differences are particularly evident for small training sets, suggesting that TransMEP performs better on small datasets, which is often a reality in protein engineering studies.

### 3.2 Wet-lab study simulation

#### Setup

In this computer experiment, we test the applicability of TransMEP for wet-lab protein engineering using the upper confidence criterion. For each *κ* value considered (see Eq. 8), 100 simulations are performed. Each of these simulation studies starts with an initial sample consisting of the wild type and nine randomly sampled mutants with a single mutation. Thereafter, the next mutant to be tested is selected iteratively according to the UCB criterion until convergence is reached. Each study is limited to a total of 100 mutants, as this number can be realistically tested in the wet lab (which is not done here). In the end, the mutant with the highest target value predicted by the final model is selected.

#### Data

To simulate wet-lab studies, two large protein mutagenesis datasets are used. The first dataset is from Wu et al. ^30^ who provided a large collection of mutants of protein G domain B1 (GB1) from *Streptococcal bacteria*. The mutations are restricted to four positions, resulting in 20^4^ = 160,000 possible variants, of which 149,361 (93%) are recorded in the dataset. The target value for these mutants is their ability to fold into the native GBP1 structure, which is indirectly measured by the binding affinity to the Fc-fragment of the immunoglobulin G antibody (IgG-Fc). This dataset contains many epistatic effects, i.e., non-linear interactions of mutations, which makes it particularly interesting for machine learning.^10^ The other dataset, provided by Sarkisyan et al.,^31^ contains 51,715 mutants of the green fluorescent protein (GFP). The mutations were created randomly, making it an unbiased dataset. As such, the dataset contains significantly more mutated positions than the GB1 dataset, but has less coverage of the complete mutation space. The fluorescence of the GFP is used as the target value.

#### Results

The results of this computer experiment are the target values of the predicted best mutants, which are shown in three plots per protein in Figure 2. Most notably, UCB optimization on TransMEP yields very good variants for both proteins. For the *κ* values that lead to the best results, TransMEP achieves a final mutant with a target mean in the upper 0.3% of all variants present in either data set. The optimal *κ* value depends on the dataset.

**Figure 2:**
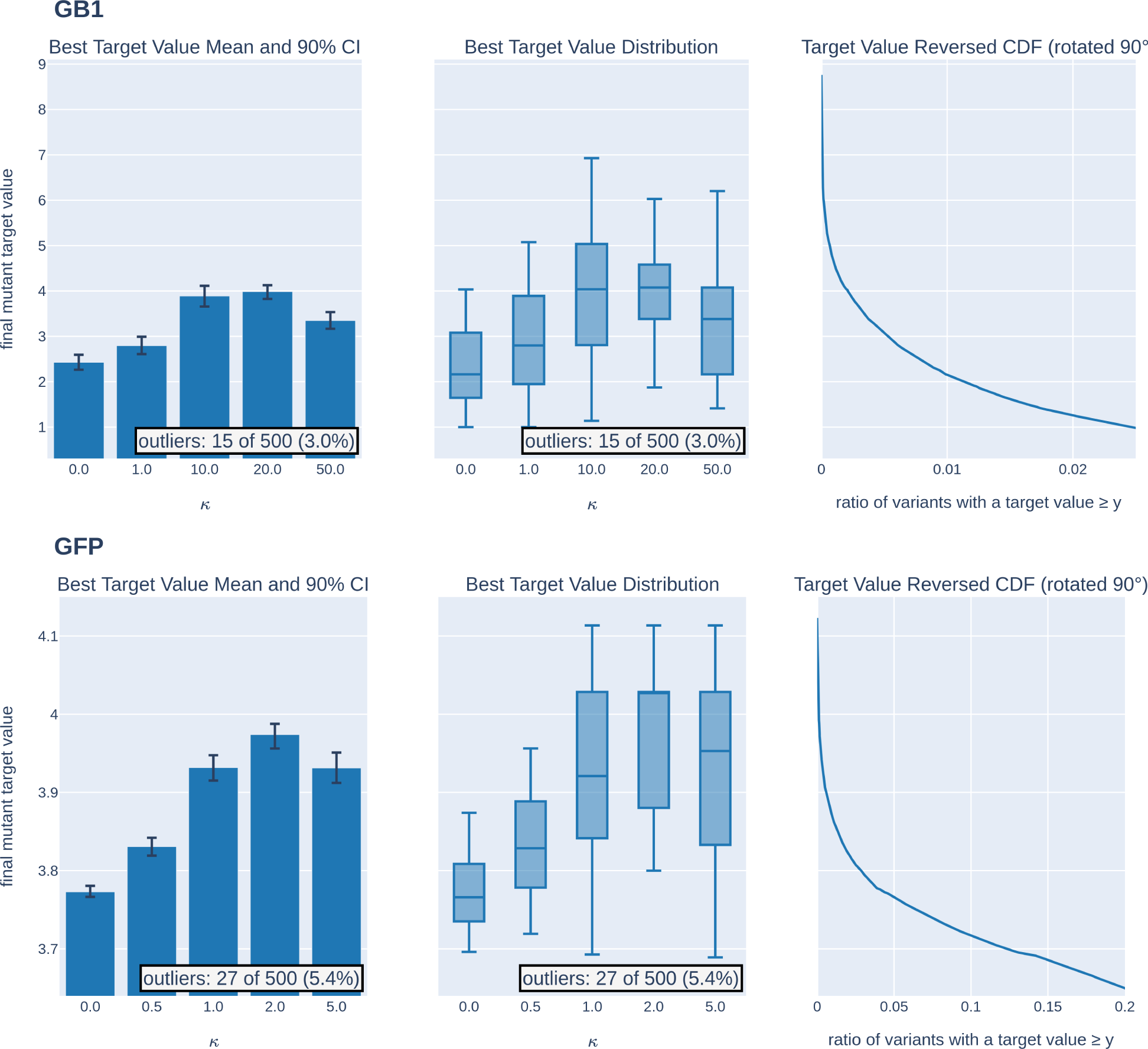
Results of the wet-lab study simulations on the datasets of GB1 (top) and GFP (bottom). (Left) The mean of the target value for the final mutant resulting from each of the 100 simulations with 90% confidence interval (CI) computed using bootstrapping is shown for increasing *κ* values. (Middle) The target value distributions are shown as boxplots. (Right) The reversed cumulative distribution function (CDF) of the target values, defined as 1 *−* CDF and rotated by 90*^◦^*, is shown. For a point (*x, y*) on this curve, *x* corresponds to the ratio of variants with at least the target value *y*. Outlier studies with a value further away than 1.5 times the interquartile range were removed.

#### Discussion

This study shows that the combination of TransMEP with UCB optimization is promising for wet-lab studies. Interestingly, pure exploitation (*κ* = 0), i.e., only using the predicted mean as an indicator, performs significantly worse than mixing exploitation with exploration (*κ >* 0), even for the small number of mutants predicted here. This suggests that the predicted standard deviation, i.e., a confidence measure provided by Gaussian process regression, is key to improving optimization performance. However, determining the optimal *κ* value for a given wet-lab study remains difficult. Automatic HPO is not applicable here as this would require multiple independent wet-lab studies, which is prohibitive in cost. Good optimization results are possible despite the bad prediction quality of TransMEP with random training sets of the same size (R^2^ *≈* 0.05 for GB1 and R^2^ *≈* 0.28 for GFP).

Therefore, the selection of interesting mutants to test in vitro is important. Note that these simulations use the fact that the search space for finding a sequence that maximizes the UCB criterion is limited to the known mutants.

### 3.3 Sampling of high UCB Values

#### Setup

As discussed, special sampling techniques are necessary to find a candidate with a high UCB value according to a given model. Here, the previously introduced genetic algorithm is compared to random search and the UCB values of all samples in the dataset. All approaches are tested on five models that were trained with 100 random training samples each and restricted to *k* = 5 mutations. Each genetic optimization is started 10 times with different initial populations and uses hyperparameters *n* = 1000, *p_c_* = 0.5, *p_m_* = <u>^1^</u> , where *m* is the sequence length (see section 2.5 for the other definitions), and a maximum of 100 iterations. Random search is also started 10 times, and it is restricted to the same computation time as genetic optimization.

#### Data

This computational experiment uses the GFP dataset introduced earlier. The GB1 dataset is not of interest here because there are only 20^4^ = 160,000 variants that can be exhaustively scanned with a modern GPU, i.e. if a user wants to apply TransMEP to this protein, they do not need genetic optimization but can follow a brute force approach.

#### Results

In general, genetic optimization finds mutants with significantly higher UCB values than random search (Figure 3). Additionally, the found mutants exhibit higher UCB values than the mutants already present in the dataset. The differences in achieved UCB values between models is high.

**Figure 3:**
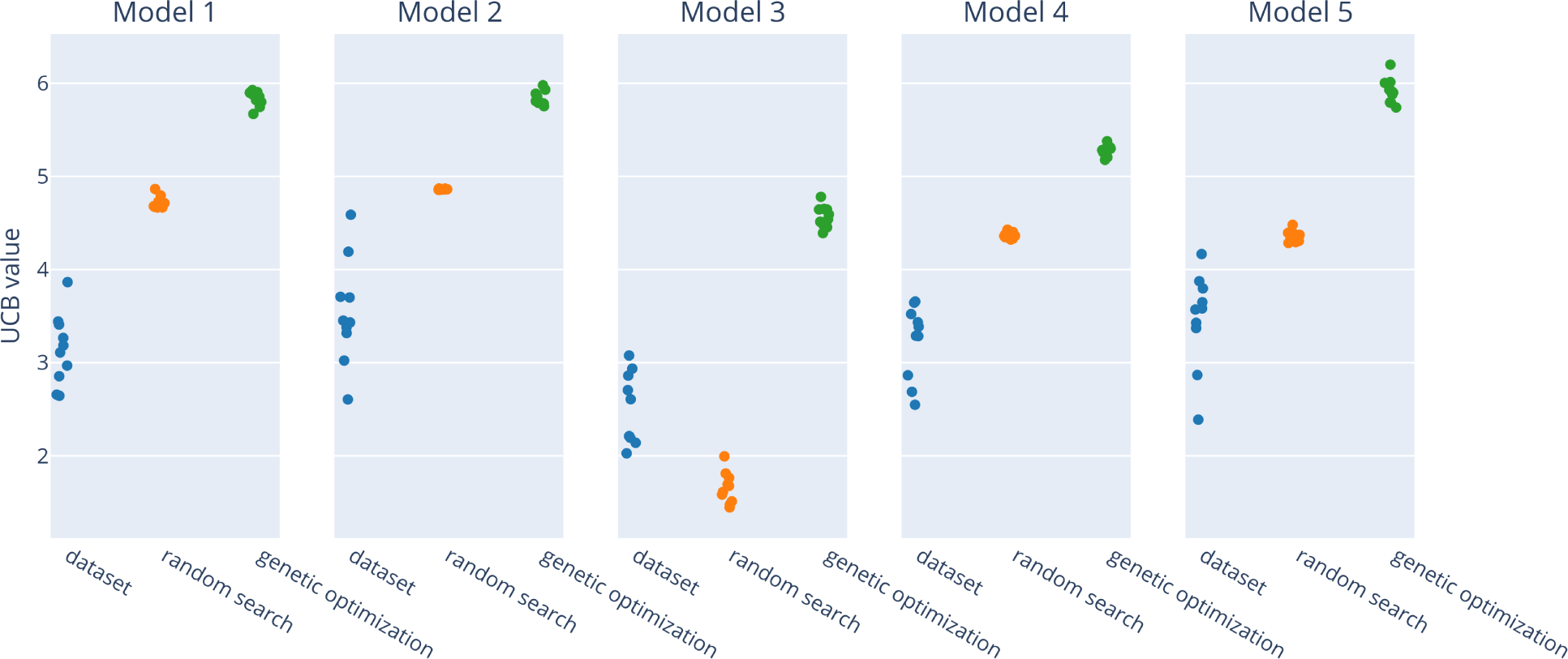
Top 10 mutants found in the dataset, by random search and genetic optimization. Each model was trained using 100 samples from the GFP dataset and random search was restricted to the computing time of genetic optimization.

#### Discussion

Most importantly, genetic optimization provides improvements over the naive approach of random search. As such, it is useful for finding mutants with high UCB values. However, the maximum achievable UCB value and thus the potential for further optimizations remains unknown. Interestingly, there are potentially more promising mutants found by genetic optimization than available in the dataset, which would be interesting to test in vitro in the future

## 4 Conclusion

In this work, transfer learning on PLMs was applied to mutation effect prediction using Gaussian process regression. This increased the prediction quality over fine-tuned PLMs while reducing runtime and improving training stability. The approach was implemented in a user friendly software tool called TransMEP available on GitHub (https://github. com/strodel-group/TransMEP). Further, wet-lab studies were simulated to assess the effectiveness of UCB optimization for protein engineering. This yielded promising results and demonstrated the importance of exploration for successful optimization. Although TransMEP does not always achieve very good R^2^ values on all datasets, it is still highly useful for optimization. As such, good optimization results do not depend on a machine learning algorithm that can predict the whole target value landscape well. However, improvements in the machine learning algorithm as in our work presumably lead to better optimization results.

There are several ways to extend this research. One possibility would be to further optimize the Gaussian process regression with more advanced kernel function formulations to improve predictive performance. Other machine learning approaches that predict confidence measures, such as other Bayesian learning approaches, could also be of use. In addition, several problems that arise in the context of UCB optimization could be addressed. For example, the algorithm could be further optimized to find the maximizer of the UCB criterion. Another challenge is to determine the *κ* value a priori. Here, an adaptive solution would be preferable, which performs more exploration the closer it gets to convergence. Finally, it may be advantageous to sample multiple mutants simultaneously for analysis of in vitro to enable the use of a batch procedure in the wet lab. This would require to switch from UCB to a batched optimization criterion that takes both exploration and exploitation into account.

## Acknowledgement

We thank Anna Jäckering and Wibke Schumann for fruitful discussions, with special thanks to Anna Jäckering for proofreading the manuscript. Our gratitude extends to the contributors to the open-source software tools used in this project. Simulations were performed with computing resources granted by RWTH Aachen University under projects rwth0794 and p0020219.

## Supporting Information Available

Links to the datasets used in this work, instructions for using the software TransMEP, approaches to the explainability of models generated by TransMEP, derivations of the optimized Gaussian process regression formulas used in the code.

