## Supplementary Methods for "TransMEP: Transfer learning on large protein language models to predict mutation effects of proteins from a small known dataset"

### 1 Datasets, software, and usage instructions

All datasets used in this study are available at the file server `https://mutation-prediction.thoffbauer.de/transmep/datasets/`. Please contact the authors if the file server is not available. The datasets are taken from publications [1–3].

The TransMEP software is available on GitHub: `https://github.com/strodel-group/TransMEP`. There, one can also find installation and usage instructions. The general workflow for using TransMEP to create optimized protein variants by combining TransMEP predictions with wet lab studies is as follows:

1. Select an initial set of mutants as a seed for future optimization. For example, one could select the wild-type protein and 9 random single-point mutations. Evaluate these mutants in the lab and collect their target values.
2. Use the obtained values and train an initial model.
3. View the TransMEP reports (note that computing $R^2$ values is not applicable in this setting).
4. Determine the next variants to test using the UCB criterion.
5. Test the predicted mutants in the lab by measuring the target value and go back to step 2. Repeat until a sufficiently high target value is achieved.

Steps 2–4 correspond to the TransMEP commands `transmep train`, `transmep reports`, `transmep predict`, and `transmep ucb`, which are explained in detail on the GitHub page.

### 2 Model explainability

TransMEP also contains procedures that support the interpretability of the model. One of these procedures allows to determine the importance of the different training sets for mutation prediction, while another procedure facilitates the determination of the effect of a single mutation on the protein variant with multiple mutations. The theoretical basis of these two procedures is described below. This is for future reference, as within the current purely theoretical study the output of these procedures could not yet be verified.

#### 2.1 Training samples importance

Considering the equation for the expected target value of an query dataset $\mathbf{x}^* \in \mathcal{X}^m$ (Equation (6) in the main article)

$$\mathbb{E}(f(\mathbf{x}^*)) = K(\mathbf{x}^*, \mathbf{x})(K(\mathbf{x}, \mathbf{x}) + \alpha I_n)^{-1}\mathbf{y} \tag{1}$$

one can observe that this can be reduced to a linear projection of the kernel matrix $K(\mathbf{x}^*, \mathbf{x})$ by a vector $\mathbf{a} \in \mathbb{R}^n$ independent of $\mathbf{x}^*$:

$$\mathbb{E}(f(\mathbf{x}^*)) = K(\mathbf{x}^*, \mathbf{x})\,\mathbf{a}, \text{ where } \mathbf{a} = (K(\mathbf{x}, \mathbf{x}) + \alpha I_n)^{-1}\mathbf{y} \tag{2}$$

As such, the prediction $p \in \mathbb{R}$ for a single sample $x^* \in \mathcal{X}$ is given by

$$p = \sum_{i=1}^{n} k(x^*, x_i)\, a_i \tag{3}$$

Therefore, one can define the importance $w_i$ of each training sample $x_i$ for the prediction of $x^*$ as

$$w_i = \frac{\tilde{w}_i}{\sum_{j=1}^{n} \tilde{w}_j}, \text{ where } \tilde{w}_i = |k(x^*, x_i)\, a_i|\,. \tag{4}$$

#### 2.2 Mutation effect

Determining the effect of a single mutation on the mutant is more involved, as it requires backpropagation through the PLM. Fortunately, both the PLM and the Gaussian Process are differentiable, and as such one can leverage existing techniques for deep learning like Integrated Gradients.[4] This approach approximates the effect of each input variable by integrating its gradient along a straight path from a baseline to the sample of interest. TransMEP uses the wild type as the baseline and integrates the gradient of each mutation until it reaches the mutant of interest. This provides an estimate of the effect of each individual mutation in the mutant of interest. The accumulated effects should sum to the total difference between the mutant and the wild type, which provides an error estimate of the approximation.

### 3 Derivation of the optimized formulas for Gaussian Process regression

The actual implementation of TransMEP optimizes the inference equations further. This section explains these derivations to improve the accessibility of the code. First, recall the original inference equation (Equation (6) in the main article):

$$\mathbb{E}(f(\mathbf{x}^*)) = K(\mathbf{x}^*, \mathbf{x})(K(\mathbf{x}, \mathbf{x}) + \alpha I_n)^{-1}\mathbf{y} \tag{5}$$

The main computational complexity and numerical instabilities are induced by the inverse of the regularized kernel matrix $(K(\mathbf{x}, \mathbf{x}) + \alpha I_n)^{-1}$. As such, one can first reformulate above equations by replacing the inverse with the solution of a linear equation system:

$$\mathbb{E}(f(\mathbf{x}^*)) = K(\mathbf{x}^*, \mathbf{x})\,\mathbf{a},\ \text{with}\ (K(\mathbf{x}, \mathbf{x}) + \alpha I_n)\,\mathbf{a} = \mathbf{y} \tag{6}$$

As $K(\mathbf{x}, \mathbf{x})$ is a kernel matrix, it is symmetric and positive definite, and thus $K(\mathbf{x}, \mathbf{x}) + \alpha I_n$ is symmetric and positive definite, too. Therefore, one can use the Cholesky decomposition $BB^T = K(\mathbf{x}, \mathbf{x}) + \alpha I_n$ with $B \in \mathbb{R}^{n\times n}$ being a lower triangle matrix. Solving the linear equation system $BB^T\mathbf{a} = \mathbf{y}$ is fast due to optimizations that can be performed given the lower triangular form of $B$. Additionally, solutions to a linear equation system are numerically more stable than a matrix inverse. The solution $\mathbf{a}$ can be precomputed at training time and cached for inference.

Moving to the prediction of the confidence, the Cholesky decomposition can be applied as well. Again, recall the original confidence inference equation (Equation (7) in the main article):

$$\text{cov}(f(\mathbf{x}^*)) = K(\mathbf{x}^*, \mathbf{x}^*) - K(\mathbf{x}^*, \mathbf{x})(K(\mathbf{x}, \mathbf{x}) + \alpha I_n)^{-1} K(\mathbf{x}, \mathbf{x}^*) \tag{7}$$

Replacing the regularized kernel matrix with its Cholesky decomposition, one obtains

$$\begin{aligned}
\text{cov}(f(\mathbf{x}^*)) &= K(\mathbf{x}^*, \mathbf{x}^*) - K(\mathbf{x}^*, \mathbf{x})(BB^T)^{-1} K(\mathbf{x}, \mathbf{x}^*) && (8)\\
&= K(\mathbf{x}^*, \mathbf{x}^*) - K(\mathbf{x}^*, \mathbf{x}) B^{-T} B^{-1} K(\mathbf{x}, \mathbf{x}^*) && (9)\\
&= K(\mathbf{x}^*, \mathbf{x}^*) - (B^{-1} K(\mathbf{x}^*, \mathbf{x})^T)^T (B^{-1} K(\mathbf{x}^*, \mathbf{x})^T) && (10)\\
&= K(\mathbf{x}^*, \mathbf{x}^*) - C^T C, \text{ with } BC = K(\mathbf{x}^*, \mathbf{x})^T. && (11)
\end{aligned}$$

Again, the solution of the linear equation system $C$ can be obtained quickly due to the triangular form of $B$ and is numerically more stable than the inverse of the regularized kernel matrix. As such, the confidence prediction can also be accelerated by precomputing the Cholesky decomposition. One can also re-use $K(\mathbf{x}^*, \mathbf{x})$ from above. By further ignoring the covariances and only computing the variances, one observes that the diagonal of the kernel matrix $K(\mathbf{x}^*, \mathbf{x}^*)$ is $\mathbf{1}$ due to the choice of an RBF kernel (cf. Equations (2) and (3) in the main article), eliminating the need to compute $K(\mathbf{x}^*, \mathbf{x}^*)$. This leads to the following equation:

$$\text{var}(f(\mathbf{x}^*)) = \mathbf{1} - \mathbf{c}, \text{ where } \mathbf{c}_i = ||C_i||_2^2 \text{ for } i \in \{1, \ldots, n\} \text{ and } BC = K(\mathbf{x}^*, \mathbf{x})^T \tag{12}$$

Here, $C_i$ denotes the $i$-th column of $C$. By accelerating confidence prediction, the evaluation of the UCB criterion for a large number of samples is accelerated significantly.
